# Quantifying the surveillance required to sustain genetic marker-based antibiotic resistance diagnostics

**DOI:** 10.1101/699918

**Authors:** Allison L. Hicks, Stephen M. Kissler, Marc Lipsitch, Yonatan H. Grad

**Affiliations:** Department of Immunology and Infectious Diseases, Harvard T. H. Chan School of Public Health, Boston, Massachusetts 02115; Center for Communicable Disease Dynamics, Department of Epidemiology, Harvard T. H. Chan School of Public Health, Boston, Massachusetts 02115; Division of Infectious Diseases, Department of Medicine, Brigham and Women’s Hospital, Harvard Medical School, Boston, Massachusetts 02115

**Author notes:** **Corresponding author**: Yonatan H. Grad, MD, PhD, Harvard T. H. Chan School of Public Health, 665 Huntington Ave, Rm 715 Building 1, Boston, MA 02115. Co-senior authors (Marc Lipsitch, Yonatan H. Grad).

## Abstract

Diagnostics that minimize the time to selection of an appropriate antibiotic treatment represent an important strategy in addressing the challenge of antimicrobial resistance (AMR). Among this class of diagnostics, the use of pathogen genotype to predict AMR phenotype has been facilitated by advances in rapid sequencing platforms. A longstanding objection to this approach, however, is that the emergence of novel resistance mechanisms will inevitably lead to a decline in the sensitivity of these diagnostics. Here, we show that while the sensitivities of some genetic markers of resistance remain stably high, sensitivities of other markers rapidly decline, as expected, due to the emergence of novel resistance variants. We then present a simple mathematical framework that defines the sampling and phenotypic testing rates needed for early detection of novel resistance variants and thus demonstrate how surveillance can help maintain the sensitivity and utility of sequence-based AMR diagnostics.

**One sentence summary:** Targeted sampling strategies are necessary for early detection of novel resistance mechanisms and sustainability of genotype-based detection of novel resistance mechanisms and sustainability of genotype-based diagnostics.

---

Antimicrobial resistance (AMR) poses a grave threat to global public health, underscoring the need for strategies to slow and control the spread of resistance. One direction is to develop fast and reliable diagnostics that minimize the delay between diagnosis and selection of an appropriate treatment regimen based on the target pathogen’s antibiotic susceptibility profile (*1, 2*). A promising approach, use of pathogen genotype to predict AMR phenotype, has been facilitated by advances in rapid and cost-efficient amplification and sequencing. For example, the Cepheid GeneXpert MTB/RIF assay for rifampicin resistance in *Mycobacterium tuberculosis*, and the SpeeDx ResistancePlus GC assay for ciprofloxacin resistance in *Neisseria gonorrhoeae*, are already in clinical use, and many others are in the pipeline (*3-5*). These diagnostics must maintain high sensitivity to remain useful clinically. However, the emergence of novel resistance mechanisms will inevitably lead to a decline in sensitivity, perhaps exacerbated by variable prevalence of resistance determinants across populations (*6*). Key to maintaining sensitivity is therefore sustained sampling and routine updating of the diagnostics with newly described resistance determinants. However, the rate of sampling necessary for timely detection of novel resistance variants has been unclear.

Here, we use datasets of clinical isolates of multiple pathogens collected over 7-14 years to show that while the sensitivities of some genetic markers of resistance remain stably high, sensitivities of other markers rapidly decline due to the emergence of novel resistance variants. We present a simple mathematical framework that defines the rates of sampling and phenotypic testing necessary for early detection of novel resistance variants.

## Waning sensitivity of resistance markers

In the ideal scenario for a genotype-based antibiotic resistance diagnostic, phenotypic resistance is always encoded by a specific genotype, e.g., a single, stereotyped mutation or gene. To date, some combinations of bacteria and antibiotics approximately satisfy this criterion: target modification mutations in *gyrA* maintain high sensitivity for predicting ciprofloxacin non-susceptibility in *N. gonorrhoeae* and *Acinetobacter baumannii*, **Fig. 1A-B**. For others, the genetic markers of resistance show decreased sensitivity over time, corresponding to increased incidence of previously rare or undetected resistance markers (**Fig. 1C-E**). The *gyrA* target modification mutation in *Klebsiella pneumoniae* isolates (*7*) (NCBI BioProjects SRP102664, SRP110988, and SRP116139), for example, becomes a less sensitive predictor of ciprofloxacin non-susceptibility as the incidence of isolates with acquired *qnr* genes (which code for target protecting proteins) increases (**Fig. 1C**). Similarly, the emergence of the mosaic *penA* (XXXIV) allele in *N. gonorrhoeae* clinical isolates (*8*) (NCBI BioProjects ERP008891, ERP001405 and ERP000144) corresponds to decreased sensitivity of other target modification mutations for predicting penicillin non-susceptibility (**Fig. 1D**). Further, decreased sensitivity of *bla*_OXA-58_ for predicting imipenem non-susceptibility in *A. baumannii* clinical isolates from the U.S. military healthcare system (NCBI BioProject SRP065910) is associated with increased incidence of other oxacillinases (**Fig. 1E**).

**Fig. 1.**
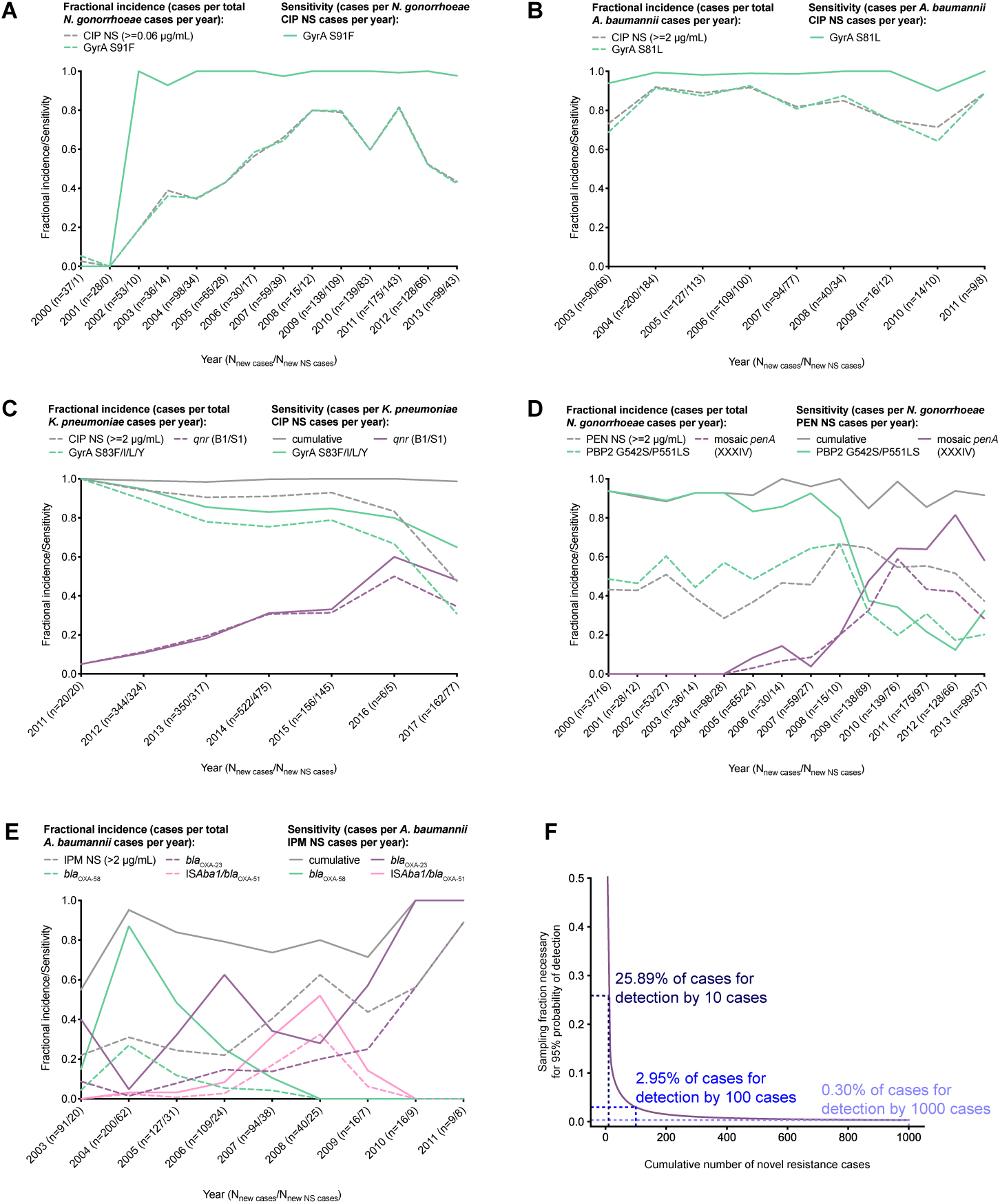
Emergence of novel resistance variants and a framework for their detection. Fractional incidence and sensitivity in predicting ciprofloxacin (CIP), penicillin (PEN), and imipenem (IPM) non-susceptibility (NS) of different genetic variants over time in *Neisseria gonorrhoeae, Acinetobacter baumannii*, and *Klebsiella pneumoniae* (**A-E**). Fractional incidence of different markers may not sum to 100% due to uncharacterized resistance markers or strains carrying multiple markers. Quantifying sampling efforts required for detection of novel variants as a function of the total number of novel resistance cases prior to detection (**F**).

## Defining required sampling rate as a function of diagnostic failure threshold

Given the possible emergence of novel resistance variants, maintenance of a genotype-based AMR diagnostic requires surveillance and phenotyping of clinical specimens predicted to be susceptible, characterization of novel resistance determinants, and subsequent updating of the diagnostic. Once a resistant strain not captured by the current diagnostic test appears in the population, there is a simple relationship between the cumulative number of such cases and the probability that at least one will be detected: if *f* is the proportion of all genotypically susceptible cases that receive confirmatory phenotypic testing, and *N* is the number of variant cases, then the probability *x* that the new variant is detected in at least one of those cases is given by *x* = 1 − (1 − *f*)^*N*^. Therefore, to have a probability of at least *x* that the new variant will be detected by the time *N* cases of it have occurred, the proportion receiving confirmatory testing must be

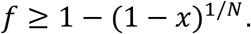

Thus, to be 95% confident that a novel variant is detected by the time it has occurred in a total of 100 cases, approximately 3% of cases must be phenotypically tested (**Fig. 1F**). Given the 555,608 cases of gonorrhea in the U.S. in 2017, (https://www.cdc.gov/std/stats17/gonorrhea.htm), the required sampling rate for detection of a novel variant by the time it occurred in 100 cases would be 1,367 cases per month. However, for surveillance programs aimed at detecting novel resistance variants that undermine the sensitivity of a genotype-based diagnostic that has already been implemented in the population, cases with isolates predicted to be resistant by the diagnostic would be excluded from the sampling population, reducing the required sampling rate.

If the variant is increasing in frequency in a population at a rate *r* per unit time, then the time (beginning at *t*_*0*_, when the variant first emerged in a single case) at which there is a probability of 1 − *x* of having detected the variant is:

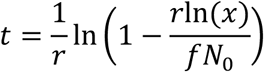

where *N*_*0*_ is the initial population-wide incidence of the variant in cases per unit time.

## Additional considerations for implementation

The practical implementation of this sampling model requires consideration of multiple additional factors. First, if the phenotyping of the collected clinical isolates is batched, then the intervals between testing could lead to delays in detection. However, the delay is bounded by the selected threshold of allowed failures and the testing interval. For example, if 3% of all genotypically susceptible strains are phenotyped, and the phenotyping is batched monthly, then the maximum delay between the occurrence of the 100th case with the novel variant and the detection of the variant will be approximately one month. Given a novel variant with a growth rate between 0.02 and 0.12 per month (as is observed for mosaic *penA* in *N. gonorrhoeae, bla*_OXA-23_ and IS*Aba1*/*bla*_OXA-51_ in *A. baumannii*, and *qnr* in *K. pneumoniae*), a maximum of between 3 and 15 additional cases of the novel variant might occur between the occurrence of the 100^th^ case and detection of the novel variant.

Second, changes in disease incidence impact the surveillance and sampling strategy. For example, gonorrhea incidence in the U.S. increased 65% between 2008 and 2017 and 18.6% between 2016 and 2017 alone (https://www.cdc.gov/std/stats17). To maintain the same level of confidence that the novel variant will be detected by the desired time or threshold number of cases, disease incidence would need to be closely monitored and surveillance matched accordingly. Given the directly proportional relationship between case incidence and sampling rate, an 18.6% increase in incidence of genotypically susceptible strains must correspond to an 18.6% increase in sampling rate in order to maintain the same confidence in detection of a novel variant by the time it has appeared a given number of cases.

Further, the incidence of clinical isolates predicted to be susceptible (susceptible case incidence), rather than overall case incidence, is of primary relevance for detecting novel resistance determinants. Thus, a third issue to consider in sampling strategy is that the susceptible case incidence may be subject to more rapid changes than overall case incidence, depending on varying selective pressures for or against resistance introduced by a variety of factors, including antibiotic use and the diagnostic itself. Thus, in establishing a plan for a sampling strategy, a conservative approach would be account for these fluctuations by calculating the necessary sampling rate as a fraction of all cases.

Relatedly, demographic and geographic heterogeneity in selective pressures, and thus in the likelihood of emergence of novel resistance variants, introduces an additional complication in selecting the populations for surveillance and sampling. Behavioral and socioeconomic factors may contribute to differential emergence of antibiotic resistance across sub-populations (*9*), and certain resistance mechanisms and variants may be more likely to appear in specific sub-populations or transmission networks *(6).* For example, for some pathogens, settings such as oncology and critical care units within hospitals, where antibiotic use is highest and where patients who have failed prior antibiotic therapies are likely to concentrate, may provide ideal locations for surveillance. Thus, while a diverse sampling of the population may be optimal in the absence of epidemiological analysis of risk factors for emergence of resistance, the latter may facilitate more targeted sampling strategies that help to reduce delays in detection of novel variants.

Delays in updating genotype-based diagnostics may also influence the rates of emergence of new variants, as these diagnostics introduce selective pressure against isolates with the diagnostic targets and increased fitness for those lacking the targets (*10-12*). Thus, assay adaptability is likely to be an important determinant of diagnostic sustainability. For genotype-diagnostics that rely on testing for specific alleles, once specimens with unknown pathways to resistance have been identified, it will be important to define the genetic basis of resistance and incorporate it into the diagnostic assay. Thus, long-term support of such diagnostics will require a system for rapidly determining the genetic basis of resistance in novel resistant variants, an activity that is currently challenging for some pathogen species with less tractable genetics. This requirement may create an advantage for diagnostics that rely on phylogenetic similarity (*13*) and are agnostic to the resistance determinant, where genetic experiments could be avoided but for which regularly updating the reference database will be critical for maintaining sensitivity.

This model is based on the assumption that the most efficient and reliable method for detection of novel resistance variants is routine phenotypic testing of strains predicted to be susceptible, but identification of treatment failures represents an additional and potentially more efficient route to detection. Depending on factors such as overall case incidence and severity of clinical failure associated with the pathogen, identification of treatment failures may be a more practical alternative to large-scale phenotypic sampling programs. However, identification of treatment failures may be encumbered by a number of factors, including long treatment regimens and/or partial abatement of symptoms and thus failure to follow up. Further, infections might be cleared even in the case of undetected resistance, and multi-drug therapy may similarly mask novel resistance to individual drugs. For example, one of the first identified cases of infection with *N. gonorrhoeae* with ceftriaxone resistance and intermediate azithromycin resistance in the UK was identified as NAAT-negative two weeks after treatment with ceftriaxone and azithromycin (*14*). Thus, while continued collection of clinical outcome data is crucial to defining the relationship between phenotypic susceptibility test results and expected treatment outcome, surveillance programs designed to regularly sample a sufficient fraction of isolates in a given population, incorporating relevant epidemiological information, may represent the most reliable strategy for comprehensive detection of novel resistance variants.

## Acknowledgements

This work was supported by Grant U54GM088558 (Models of Infectious Disease Agent Study, Center for Communicable Disease Dynamics) from the National Institute of General Medical Sciences and Grant R01AI132606 from the National Institute of Allergy and Infectious Diseases. The content is solely the responsibility of the authors and does not necessarily represent the official views of the National Institute of General Medical Sciences, National Institute of Allergy and Infectious Diseases, or National Institutes of Health.

